# Decoding Multivoxel Representations of Affective Scenes in Retinotopic Visual Cortex

**DOI:** 10.1101/2020.08.06.239764

**Authors:** Ke Bo, Siyang Yin, Yuelu Liu, Zhenhong Hu, Sreenivasan Meyyapan, Sungkean Kim, Andreas Keil, Mingzhou Ding

**Author notes:** **Correspondence:** Mingzhou Ding and Andreas Keil.

## Abstract

The perception of opportunities and threats in complex scenes represents one of the main functions of the human visual system. In the laboratory, its neurophysiological basis is often studied by having observers view pictures varying in affective content. This body of work has consistently shown that viewing emotionally engaging, compared to neutral, pictures (1) heightens blood flow in limbic structures and frontoparietal cortex, as well as in anterior ventral and dorsal visual cortex, and (2) prompts an increase in the late positive event-related potential (LPP), a scalp-recorded and time-sensitive index of engagement within the network of aforementioned neural structures. The role of retinotopic visual cortex in this process has, however, been contentious, with competing theoretical notions predicting the presence versus absence of emotion-specific signals in retinotopic visual areas. The present study used multimodal neuroimaging and machine learning to address this question by examining the large-scale neural representations of affective pictures. Recording EEG and fMRI simultaneously while observers viewed pleasant, unpleasant, and neutral affective pictures, and applying multivariate pattern analysis to single-trial BOLD activities in retinotopic visual cortex, we identified three robust findings: First, unpleasant-versus-neutral decoding accuracy, as well as pleasant-versus-neutral decoding accuracy, were well above chance level in all retinotopic visual areas, including primary visual cortex. Second, the decoding accuracy in ventral visual cortex, but not in early visual cortex or dorsal visual cortex, was significantly correlated with LPP amplitude. Third, effective connectivity from amygdala to ventral visual cortex predicted unpleasant-versus-neutral decoding accuracy, and effective connectivity from ventral frontal cortex to ventral visual cortex predicted pleasant-versus-neutral decoding accuracy. These results suggest that affective pictures evoked valence-specific multivoxel neural representations in retinotopic visual cortex and that these multivoxel representations were influenced by reentry signals from limbic and frontal brain regions.

## Introduction

Visual media are a major source of information as well as entertainment, and as such, have become a central element in people’s lives. Using pictures or videos of varying emotional content to elicit strong viewer response is a key aspect of visual media usage. Emotionally engaging visual stimuli are viewed longer (Bradley et al., 2001), rated as being more arousing (Lang et al., 1993), and accompanied by heightened autonomic as well as neurophysiological responses (Bradley, 2009), compared to affectively neutral visual stimuli. Because of these properties, viewing affective pictures has been extensively used as a laboratory model of emotional engagement, resulting in an immense literature (Bradley et al., 2012), which includes studies aiming to characterize the neurophysiological basis of emotional picture perception (Frank & Sabatinelli, 2017; Sabatinelli et al., 2011).

Functional magnetic resonance imaging (fMRI) studies have shown that viewing emotionally arousing picture, relative to neutral pictures, prompts higher blood oxygen level-dependent (BOLD) activity in a widespread network of regions, including the amygdaloid complex, pulvinar, medial prefrontal cortex, orbitofrontal cortex, and widespread extrastriate parieto-occipital and temporal cortices, with strong responses in higher-order, but not retinotopic, visual areas (Lang et al., 1998a; Bradley et al., 2003; Norris et al., 2004; Sabatinelli et al., 2005; Bradley et al., 2015). Over the past decade, however, evidence from other measurement modalities (e.g., EEG and ERP; Thigpen et al., 2017), ranging from experimental animals (Li et al., 2019) to computational modeling (Kragel et al., 2019), has supported the strong involvement of early, retinotopic visual areas in representing emotional content. Although these findings are in line with behavioral studies showing that emotion facilitates early visual processing (Phelps et al., 2006), hemodynamic imaging work has to date failed to observe consistent differential activation between emotional and neutral pictures in primary visual cortex and other retinotopic areas (Lang et al 1998; Sabatinelli et al., 2014).

The lack of consistent findings may be partly attributable to the limitations of univariate analysis methods used in most of the prior fMRI studies. The recent advent of multivariate pattern analysis (MVPA) provides a potential avenue to close the gap. MVPA examines voxel-level activation within a region of interest (ROI) as a multivariate pattern and yields a decoding accuracy to quantify the difference between patterns of different classes of stimuli at the single subject level (Norman et al., 2006). This technique has been successfully applied in affective neuroscience and has extended the field beyond univariate studies by decoding multivoxel neural representations of emotion within specific brain regions (Ethofer et al., 2009; Peelen et al.,2010) and within largescale neural networks (Baucom et al., 2012;Saarimäki et al., 2015; Bush et al., 2017). Building on this body of work, the present study applied MVPA to systematically define the multivoxel patterns evoked by emotional stimuli within specific regions along the retinotopic visual hierarchy, extending from primary visual cortex (V1) to intraparietal cortex along the dorsal pathway, and to parahippocampal cortex along the ventral pathway, and tested the hypothesis that voxel patterns in retinotopic cortex code for emotional content.

If visuocortical activation patterns encode emotional content, this would raise the question regarding the origin of these patterns. Two mutually non-exclusive groups of hypotheses have been proposed to account for emotion-specific modulations of activity in retinotopic visual areas: First, the so-called reentry hypothesis states that the increased visual activation evoked by affective pictures results from reentrant feedback, meaning that signals arising in subcortical emotion-processing structures such as the amygdala propagate to visual cortex to facilitate the processing of motivationally salient stimuli (Lang and Bradley, 2010; Sabatinelli et al., 2005). According to tracer studies in macaques, such reentrant projections exist, and they are more sparse for retinotopic, compared to anterior visual regions (Amaral et al., 2003; Freese and Amaral, 2005), consistent with findings from univariate fMRI studies (Sabatinelli et al., 2011) and from ERP studies (Keil et al., 2002). Neuroimaging studies have also lent support to the reentry hypothesis by comparing enhanced hemodynamic responses evoked by emotionally arousing stimuli in the amygdala and in the visual cortex (Lane et al., 1997; Sato et al.,2004; Sabatinelli et al., 2005). For example, a fast-sample fMRI study demonstrated that the response enhancement in the amygdala precedes that in extrastriate visual cortex (Sabatinelli et al., 2009). Moreover, amygdala and visual cortical activity strongly co-vary during emotional picture viewing (Sabatinelli et al., 2005; Kang et al., 2016). Critically, patients with amygdala lesions showed no enhancement in visual cortical activity when fearful faces were presented even though their visual system remains intact (Vuilleumier et al., 2004). This evidence suggests that amygdala is possibly a source of the reentrant signals.

A second group of hypotheses states that sensory cortex, including retinotopic areas, may itself code for emotional qualities of a stimulus, without the necessity for recurrent processing (see Miskovic & Anderson, 2018, for a review). Evidence supporting this hypothesis comes from empirical studies in experimental animals (Li et al., 2019; Weinberger, 2004) and human observers (Thigpen et al., 2017), in which the extensive pairing of simple sensory cues such as tilted lines or sinusoidal gratings with emotionally relevant outcomes shapes early sensory responses (Miskovic & Keil, 2012). Beyond simple cues, recent computational work using deep neural networks has also suggested that visual cortex can intrinsically represent emotional value as contained in complex visual media such as video clips of varying affective content (Kragel et al., 2019).

The notions discussed above may be aligned under the perspective that novel, complex emotional scenes initially prompt widespread recurrent processing, including between retinotopic and limbic/frontal areas, prompting emotion-specific signaling in visual cortex (Bradley et al., 2012). If repeated extensively, critical stimulus properties of emotional stimuli may well be represented natively in retinotopic visual areas (McTeague et al., 2015). Hampering advancement of either group of hypotheses, however, is the fact that it is unclear to what extent retinotopic visual areas contain information specific to emotional content and—if so—how this information emerges. Here, we use multimodal imaging together with novel computational techniques to examine the hypotheses that (1) retinotopic visual cortex contains voxel pattern information that is specific to emotional content, and (2) these patterns are formed under the influence of reentrant signals from anterior brain structures, e.g., the amygdala.

Recurrent processing such as the engagement of large-scale reentrant feedback projections may be assessed in different ways. First, slow hemodynamic inter-area interactions can be quantified through suitable BOLD-based connectivity analyses, allowing us to quantify for example the functional connectivity between amygdala and visual cortex (Freese and Amaral, 2005; Sabatinelli et al., 2009; Sabatinelli et al., 2014). Second, recurrent processing may be characterized by leveraging the greater time resolution of scalp-recorded brain electric activity. For example, human EEG studies have shown that the late positive potential (LPP), a positive-going, long-lasting ERP component that starts about 300 – 400 ms after picture onset, may serve as an index of signal reentry. Robust LPP enhancement has been found when comparing emotion stimuli with neutral stimuli (Cacioppo et al., 1994; Schupp. et al., 2000; Keil et al., 2002; Pastor et al., 2008). Moreover, the amplitude of the LPP enhancement is linearly related to BOLD activity both in visual cortex and in the amygdala (Sabatinelli et al., 2009; Liu et al., 2012; Sabatinelli et al., 2013), and is thought to reflect heightened processing of motivational relevant stimuli in perceptual, memory, and motor systems, associated with signal reentry (Vuilleumier, 2005; Hajcak et al., 2006; Lang and Bradley, 2010).

In the present study simultaneous EEG-fMRI data were recorded from subjects viewing pleasant, unpleasant, and neutral pictures selected from the International Affective Picture System (IAPS; Lang et al., 1997) as shown in Figure 1. In addition to conventional univariate BOLD activation and ERP analysis, BOLD responses were estimated on a trial by trial basis, and MVPA was applied to single-trial BOLD responses to investigate if the neural patterns within visual cortex were distinct between emotional and neutral scenes. In addition, LPP and frontotemporal-visual cortex effective connectivity was computed as indices of reentrant feedback and correlated with MVPA decoding accuracy in visual cortex to test the impact of reentry on neural representations of emotional stimuli.

**Figure 1.**
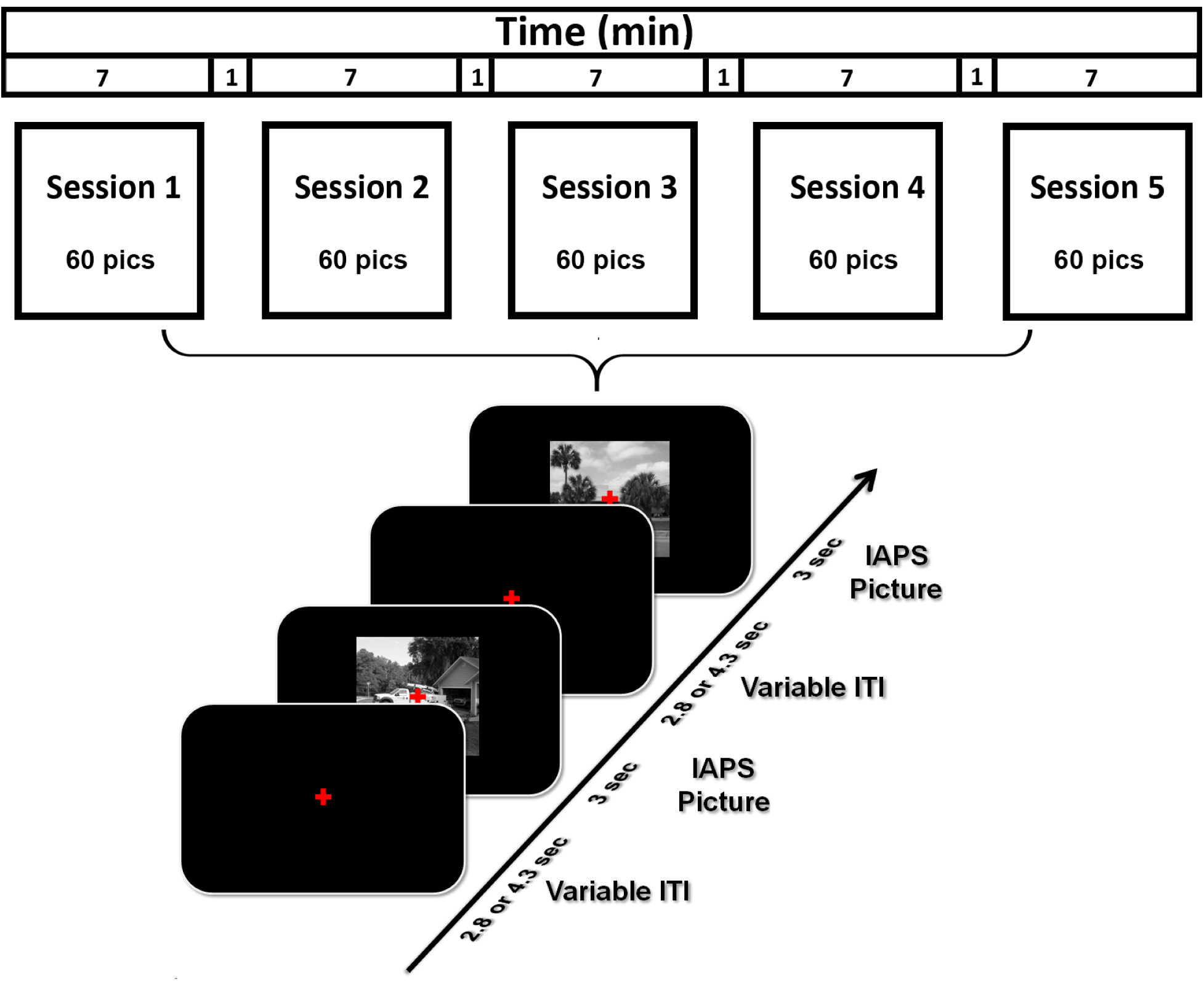
Picture viewing paradigm. There were five sessions. Each session lasted seven minutes. 60 IAPS pictures, including 20 pleasant, 20 unpleasant and 20 neutral, were presented in each session, the order of which randomly varied from session to session. Each picture lasted 3 seconds and was followed by a fixation period (2.8 or 4.3 seconds). Participants were required to fixate on the red cross in the center of screen throughout the session.

## Results

Simultaneously EEG-fMRI was recorded from human subjects viewing pleasant, unpleasant, and neutral pictures from the IAPS library. Following signal preprocessing, univariate and multivariate analysis were applied to examine the neural representations of affective pictures in retinotopic visual cortex. Because we used abbreviations quite extensively, to help with reading, we included a table in the Supplementary Materials (Table S1) where the full expression for each abbreviation is provided.

### Univariate analysis of fMRI BOLD

Relative to neutral pictures, unpleasant pictures activated bilateral occipitotemporal junctions, pre/postcentral gyrus, bilateral ventral lateral prefrontal cortices, left orbital frontal cortex, bilateral amygdalae/hippocampi, and insula (Figure 2A). Other activated areas included bilateral posterior parietal cortices, fusiform gyrus, lingual gyrus, and temporal pole. Pleasant pictures, relative to neutral, activated bilateral occipitotemporal junctions, bilateral posterior parietal cortices, right amygdala/hippocampus, bilateral inferior frontal gyrus, medial prefrontal cortex, and left orbital frontal cortex (Figure 2B). Other activated areas included fusiform gyrus, lingual gyrus, middle frontal gyrus, and temporal pole. Thus, in addition to limbic and frontal emotion processing structures, both pleasant and unpleasant affective scenes more strongly engaged regions of the higher-order visual cortex, consistent with previous reports (Lane et al., 1997; Phan et al., 2002; Liu et al., 2012; see the review by Sabatinelli et al., 2011).

**Figure 2.**
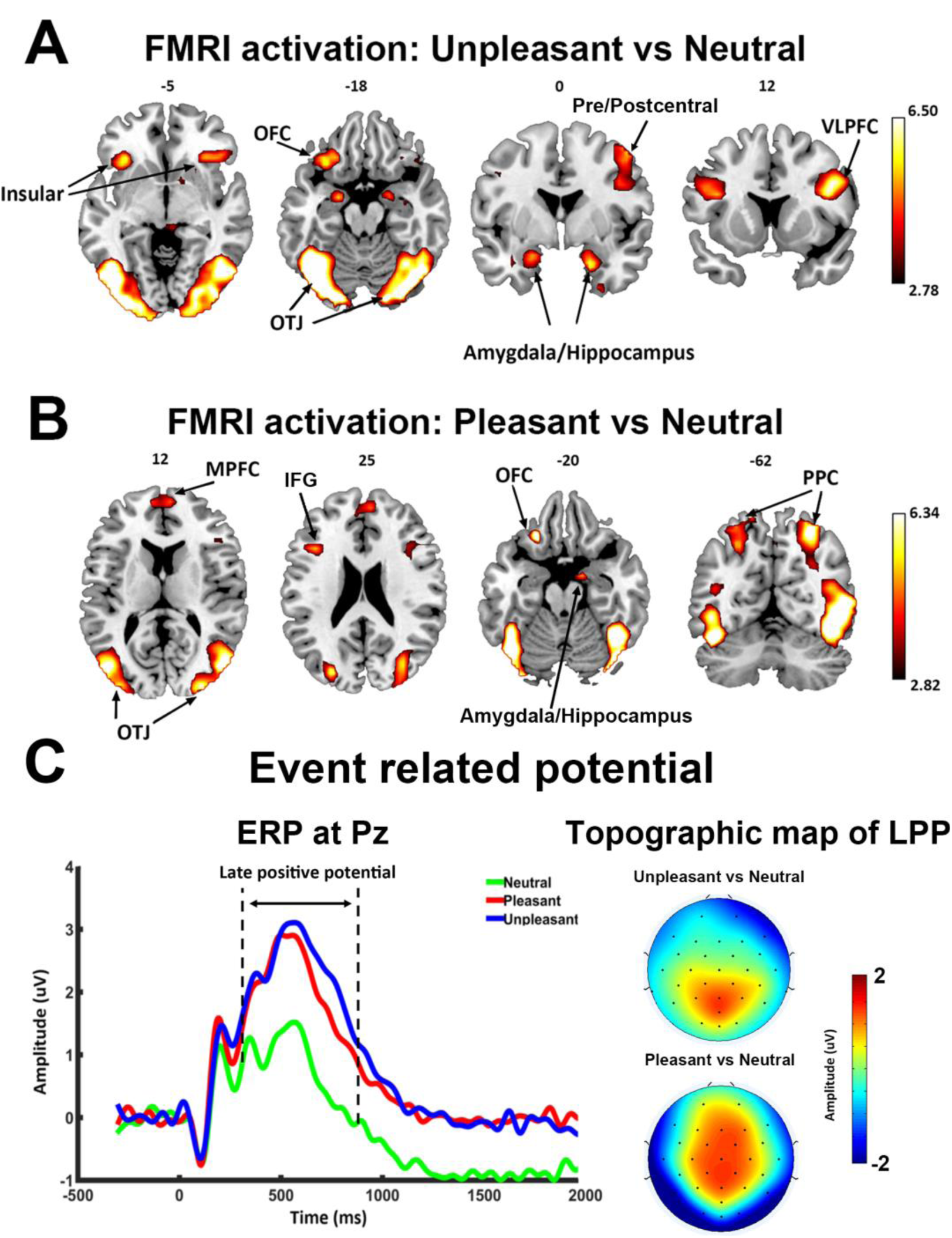
Univariate fMRI and ERP analysis. ***A***. Activation map (p<0.05, FDR) contrasting unpleasant versus neutral pictures. ***B***. Activation map (p<0.05, FDR) contrasting pleasant versus neutral pictures. ***C.*** Grand average ERP (n=20) at Pz showing ERP evoked by three classes of pictures (left) and scalp topography of LPP (300-800ms after picture onset). PPC, Posterior parietal cortex; OFC, orbital frontal cortex; MPFC, medial prefrontal cortex; OTJ, occipitotemporal junction; VLPFC, ventral lateral prefrontal cortex; IFG, inferior frontal gyrus.

#### ERP analysis

Grand average ERPs at Pz are shown for each of the three categories of pictures in Figure 2C. The late positive potential (LPP), starting ~300ms after picture onset, was higher for pleasant and unpleasant pictures compared to neutral pictures. A one-way ANOVA confirmed that LPP amplitude was significantly different among the three conditions (F=21.96, p<0.05). Post-hoc analysis confirmed that the mean LPP amplitudes for both pleasant (1.972 ± 1.882 μV) and unpleasant (2.263 ± 2.052 μV) pictures were significantly larger than that for the neutral pictures (0.781 ± 1.860 μV; pleasant vs neutral: t(19) = 4.41, p<0.001; unpleasant vs neutral: t(19) =6.52, p<0.001); no significant difference was found in LPP amplitudes between the pleasant and unpleasant categories (t(19)=1.39; p=0.18). The topographical maps depicting the scalp distribution of differential LPP amplitudes are also shown in Figure 2C. No clear differences were seen between pleasant and unpleasant pictures. These ERP results are consistent with previous reports using the same paradigm (Cuthbert et al., 2000; Keil et al., 2001; Schupp et al., 2003; Liu et al., 2012).

### MVPA analysis of fMRI BOLD

Univariate fMRI analysis in Figures 2A and 2B showed that higher-order visual cortex (e.g., fusiform, OTJ, etc.) is more strongly activated by emotional pictures, but retinotopic visual areas, including early visual cortex, are absent in the activation maps. Next, a multivariate pattern classification analysis was applied to examine the multivoxel voxel patterns evoked by the three classes of the pictures within the visual hierarchy. Retinotopic ROIs were selected according to a recently published probabilistic atlas (Wang et al., 2014), including V1d, V1v, V2d, V2v, V3v, V3d, hV4, VO1, VO2, PHC1, PHC2, hMT, LO1, LO2, V3a, V3b and IPS; see Figure 3A. The accuracy of decoding between unpleasant and neutral and between pleasant and neutral was shown for each ROI in Figure 3B. Across all visual ROIs, both unpleasant vs neutral decoding as well as pleasant vs neutral decoding were well above chance level, as determined by group level random permutation test (54% being the statistical threshold at p<0.001), demonstrating the presence of emotional signals in retinotopic visual areas including V1.

**Figure 3.**
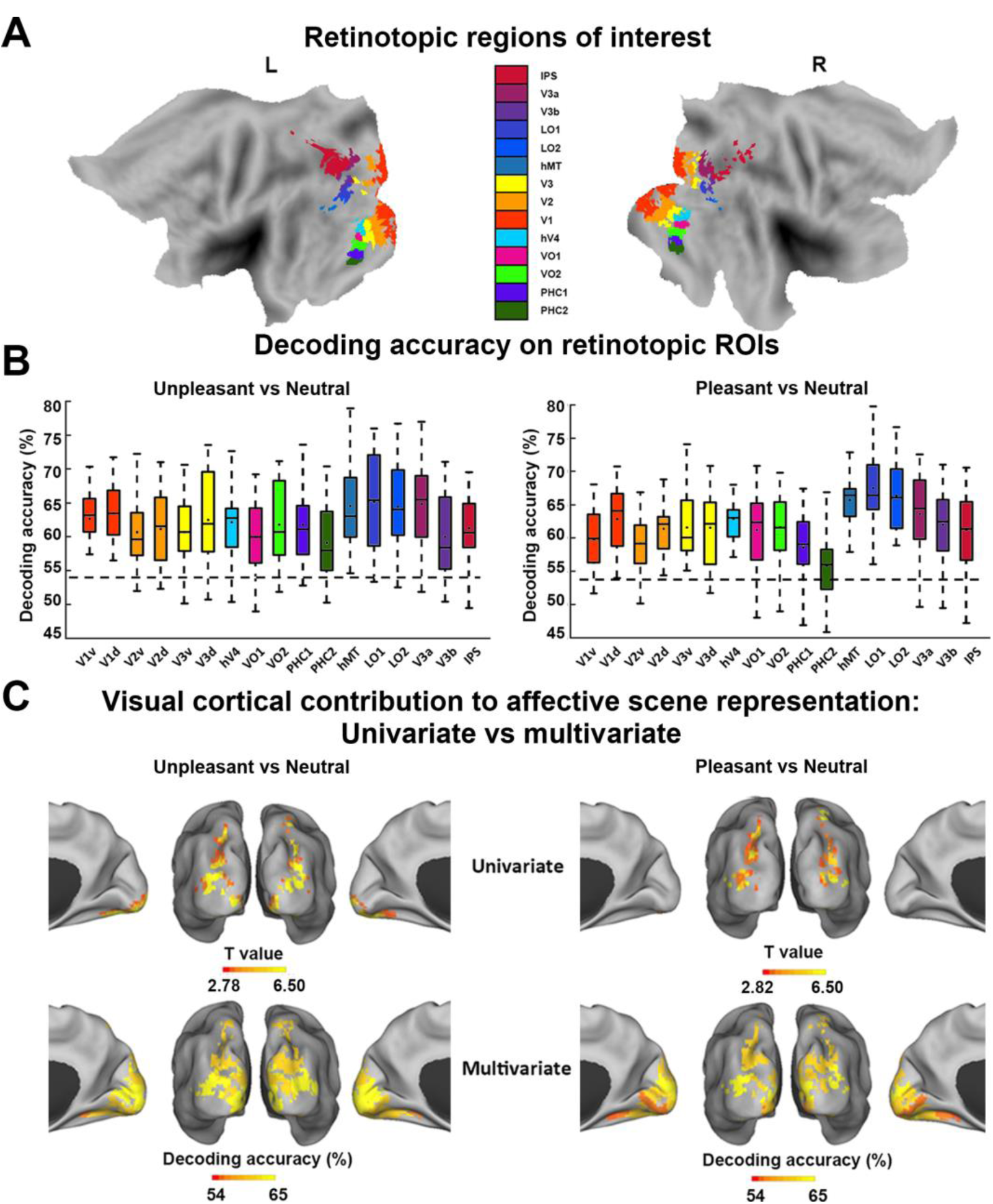
MVPA decoding analysis of neural representations of emotional scenes. ***A.*** Retinotopic ROIs visualized on the flattened brain. ***B.*** Group-average decoding accuracy between unpleasant vs neutral and pleasant vs neutral in different ROIs. Dashed line indicates the statistically significant threshold (54%). ***C***. Comparison between visual cortical contribution to the representation of affective scenes revealed by (top) univariate activation analysis (data from Figures 2A and 2B replotted) and by (bottom) multivariate decoding from Figure 3C. PHC, parahippocampal cortex. VO, ventral occipital cortex. LO, lateral occipital cortex. IPS, intraparietal sulcus. hMT, human middle temporal visual area.

Univariate analysis above identified 6 ROIs (V3d, hMT, LO1,LO2, V3a, IPS) as being more activated in pleasant vs neutral comparison (p<0.05, FDR corrected), and 11 ROIs (V2v, V3v, V3d, hV4, VO1, VO2, LO1, LO2,hMT, V3a, IPS) as being more activated in unpleasant vs neutral comparison (p<0.05, FDR corrected). Multivariate analysis, in contrast, revealed that decoding accuracy in all 17 retinotopic visual ROIs were significantly above chance level for both unpleasant-vs-neutral decoding and pleasant-vs-neutral decoding, highlighting the importance of multivoxel level activity patterns in revealing the full extent of emotional signaling in the retinotopic visual cortex (see Figure 3C for visualization).

Affective pictures are often characterized along two dimensions: valence and arousal. Relative to neutral pictures, pleasant and unpleasant pictures are different both in terms of valence as well as in terms of arousal. We attempted to control for one of the two factors (i.e., arousal) by decoding between unpleasant and pleasant pictures. As shown in Figure 4A, there was no significant difference in self-reported emotional arousal between pleasant and unpleasant pictures (p=0.2, Figure 4A), but the valence between them was significantly different (p<0.0001, Figure 4A). All the retinotopic ROIs significantly decoded the two classes of pictures above chance level (Figure 4B where 54% is the statistical threshold at p<0.001 according to the permutation test), suggesting that the differences in multivoxel patterns between pleasant vs neutral and unpleasant vs neutral comparison were simply not driven by emotional intensity of the stimuli.

**Figure 4.**
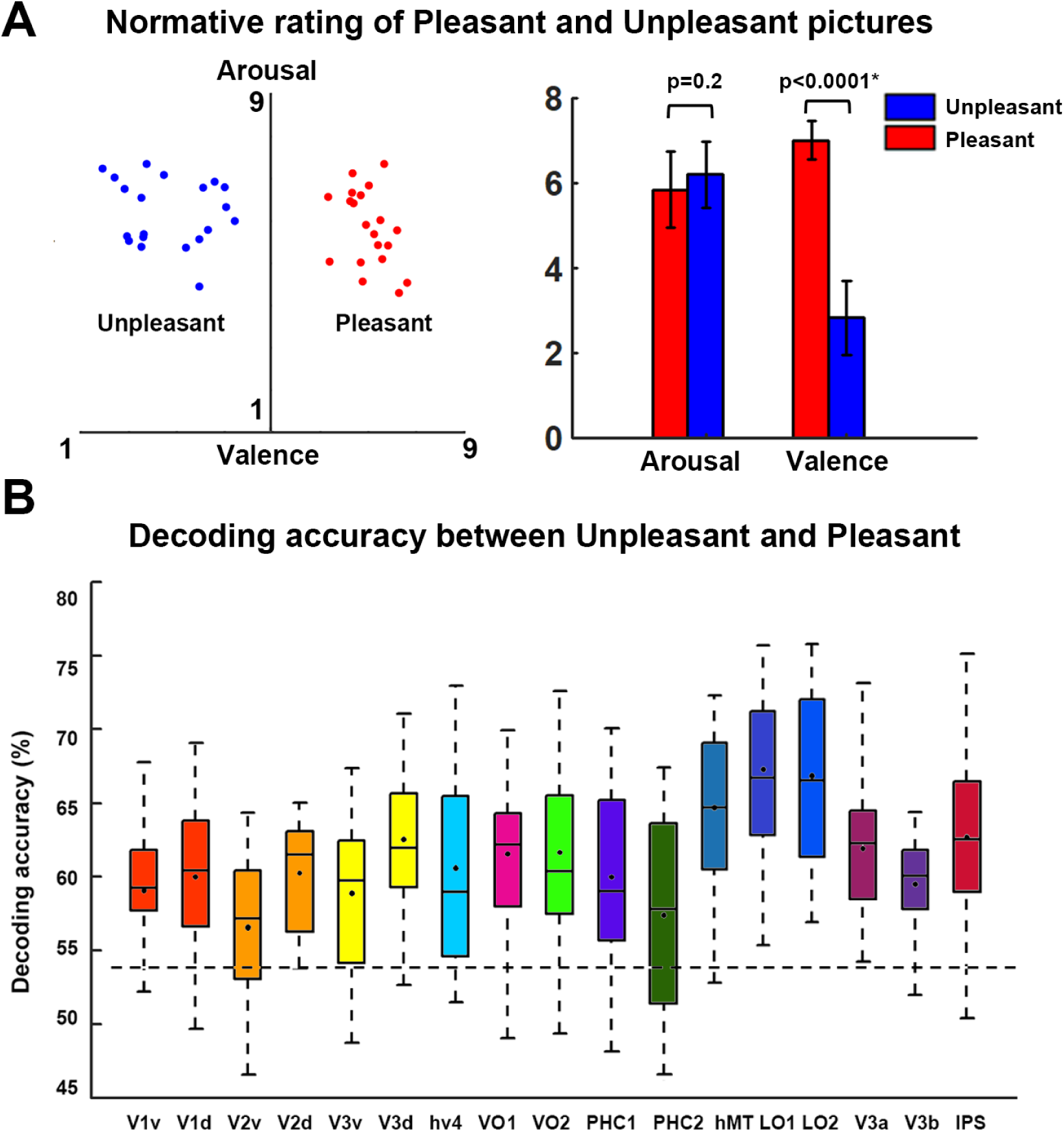
MVPA decoding of pleasant vs unpleasant scenes. ***A.*** Valence and arousal of unpleasant and pleasant images used in this study. Arousal is not significantly different between the two classes of pictures whereas pleasant pictures have significantly higher valence than unpleasant pictures. ***B.*** Group average decoding accuracy between unpleasant and pleasant in retinotopic ROIs. Dashed line indicates the statistically significant threshold (54%).

### Reentrant modulation of visual representations of affective pictures

It has been well established that viewing emotionally engaging scenes is associated with facilitated visuocortical processing, reflected in EEG, fMRI, and behavioral data. This facilitated processing is thought to be the consequence of reentrant signals that arise from emotion-modulated deep brain structures such as the amygdala and back-project into the visual system via ventral visual cortex to modulate visual processing. This hypothesis would be supported and extended to the domain of multivoxel neural representations if decoding accuracy in retinotopic areas is parametrically related to evidence of recurrent signaling. To examine this issue, we divided the retinotopic ROIs into three broader visual regions based on consistency of anatomical location and functional role: (1) early visual cortex (EVC), located on the posterior portion of the occipital lobe and involved in perception of basic visual features (Sereno et al., 1995; DeYoe et al., 1996 and Engel et al. 1997), including v1v, v1d, v2v, v2d, v3v, v3d; (2) dorsal visual cortex (DVC), located along dorsal parietal pathway and known to carry out motor and high order spatial functions such as motion perception, spatial attention, and motor preparation (Bressler et al., 2008; Konen and Kastner, 2008; Wandell and Winawer., 2011), including the combined ROIs of IPS; and (3) ventral visual cortex (VVC), located along ventral temporal pathway and known to be involved in object and scene recognition (Brewer et al. 2005; Arcaro et al. 2009), including VO1, VO2, PHC1, and PHC2. The visualized anatomical location of the three broader visual regions was shown in Figure 5A. We expect that VVC, anatomically shown as the visual structure that receives major feedback projections from the medial temporal and frontal regions (Amaral et al., 2003; Vuilleumier et al., 2004; Freese and Amara, 2005), would show significant correlation between decoding accuracy and the strength of signal reentry.

**Figure 5.**
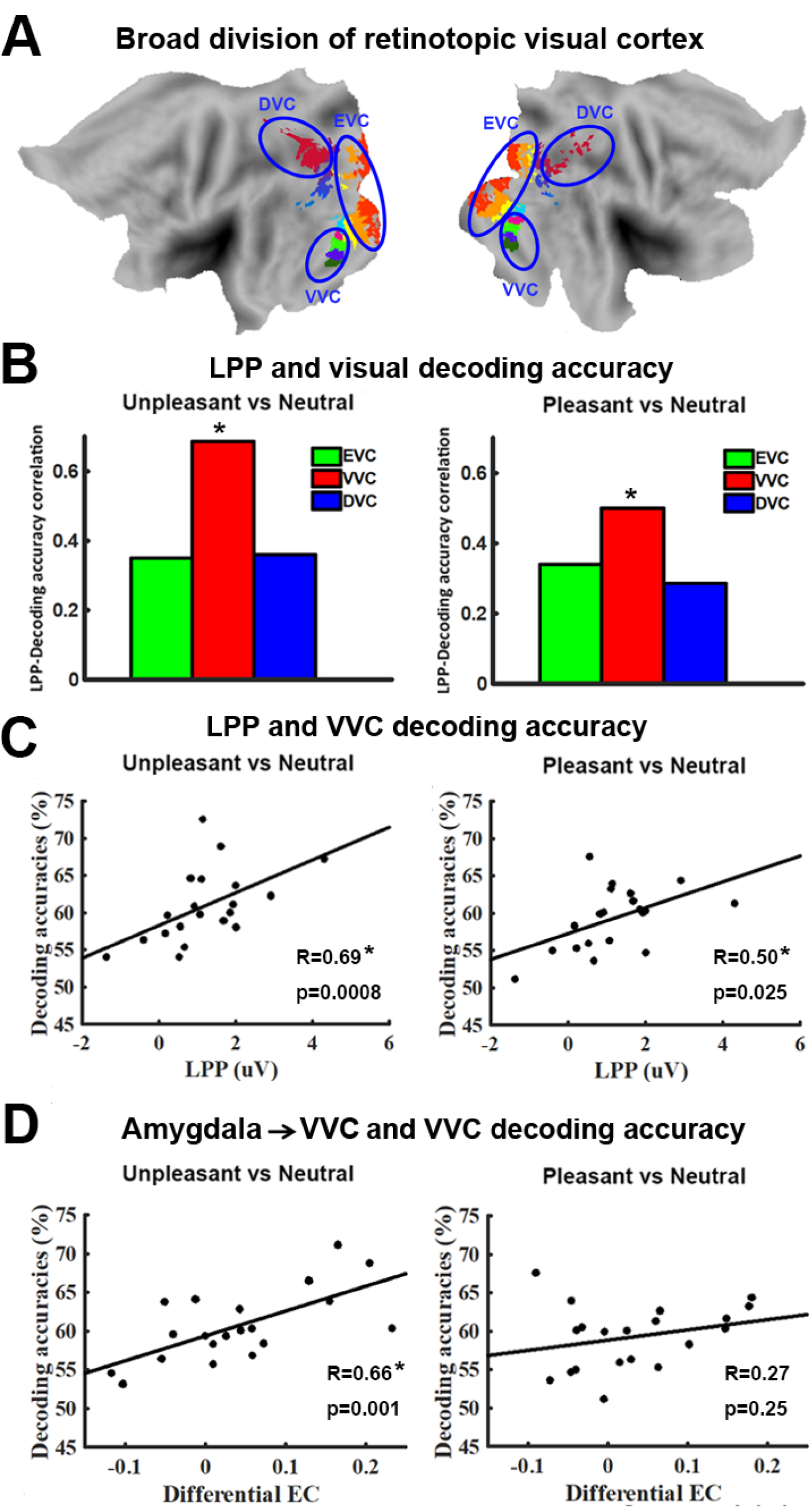
Relation between decoding accuracy and measures of signal reentry. ***A***. Anatomical location of EVC, VVC, and DVC on flattened brain. ***B***. LPP-decoding accuracy for unpleasant vs neutral (left) and pleasant vs neutral (right) in EVC, VVC, and DVC. Only decoding accuracy in VVC is significantly correlated with LPP. ***C***. Scatter plots showing relationship between LPP and decoding accuracy for unpleasant vs neutral (left) and pleasant vs neutral (right) in VVC. ***D.*** Relationship between amygdala→VVC effective connectivity and decoding accuracy in VVC. This relationship is only significant in unpleasant vs neutral comparison.

First, using LPP as an index of signal reentry, we correlated the decoding accuracy in EVC, VVC, and DVC with LPP. As shown in Figure 5B, for VVC, unpleasant vs neutral decoding accuracy and LPP amplitude were significantly correlated (R=0.69; p=0.0008); pleasant vs neutral decoding accuracy in VVC was also significantly correlated with LPP amplitude, albeit with a smaller R value (R=0.50; p=0.026). LPP was not significantly correlated with unpleasant versus neutral decoding accuracy in either EVC (R=0.37, p=0.11) or DVC (R=0.36, p=0.12). Similarly, LPP was not significantly correlated with pleasant versus neutral decoding accuracy in either EVC (R=0.34, p=0.14) or DVC (R=0.29, p=0.22).

Next, more directly assessing the amygdala to VVC feedback via DAG, a measure of directed connectivity, we computed amygdala→VVC effective connectivity, and correlated it with the decoding accuracy in VVC. Amygdala→VVC effective connectivity and VVC unpleasant versus neutral decoding accuracy was significantly correlated (r=0.66, p=0.001) (Figure 5D left), whereas amygdala→VVC effective connectivity and VVC pleasant versus neutral decoding accuracy was not significantly correlated (r=0.27; p=0.25) (Figure 5D right).

The foregoing suggested that whereas the amygdala is a likely source of reentrant signals for unpleasant picture processing, it is unlikely to be a source of reentrant signals in the processing of pleasant pictures. We then conducted a whole brain effective connectivity analysis using VVC as the seed. Effective connectivity from each voxel in the brain to VVC was computed and correlated with VVC decoding accuracy. Voxels with correlation coefficient exceeding 0.61 (p<0.005) and being part of a contiguous cluster of more than 10 such voxels were considered significant. For pleasant versus neutral comparison, as shown in Figure 6A, the potential sources of reentrant signals are bilateral STS/STG regions, right IFG, and right VLPFC, whereas for unpleasant versus neutral comparison, the potential sources of reentry are the amygdala, shown above, and the right STS/STG region (Figure 6B). To examine the collective contribution of the reentry from these regions to the neural representation of emotional pictures in ventral visual cortex, we constructed a multiple regression model using VVC decoding accuracy as the predicted variable and the following effective connectivities (EC) as predictor variables: Pleasant: IFG→VVC, VLPFC→VVC, and STS/STG→VVC; Unpleasant: Amygdala→VVC and STS/STG→VVC. Mathematically, for Pleasant, the model can be written as: *VVC decoding accuracy* = *β*_0_ + *β*_1_(*IFG* → *VVC*) + *β*_2_(*VLPFC* → *VVC*) + *β*_3_(*STS* / *STG* → *VVC*) + ε; for Unpleasant, the model can be written as:. As shown in Figure 6, the predicted decoding accuracy is strongly correlated with actual *VVC* decoding accuracy (R=0.95 for pleasant and R=0.84 for unpleasant, p<0.0001; see Figure 6A and Figure 6B respectively), demonstrating that a large portion of the variance in VVC decoding accuracy can be explained by effective connectivity from anterior temporal and prefrontal regions back to ventral visual cortex.

**Figure 6.**
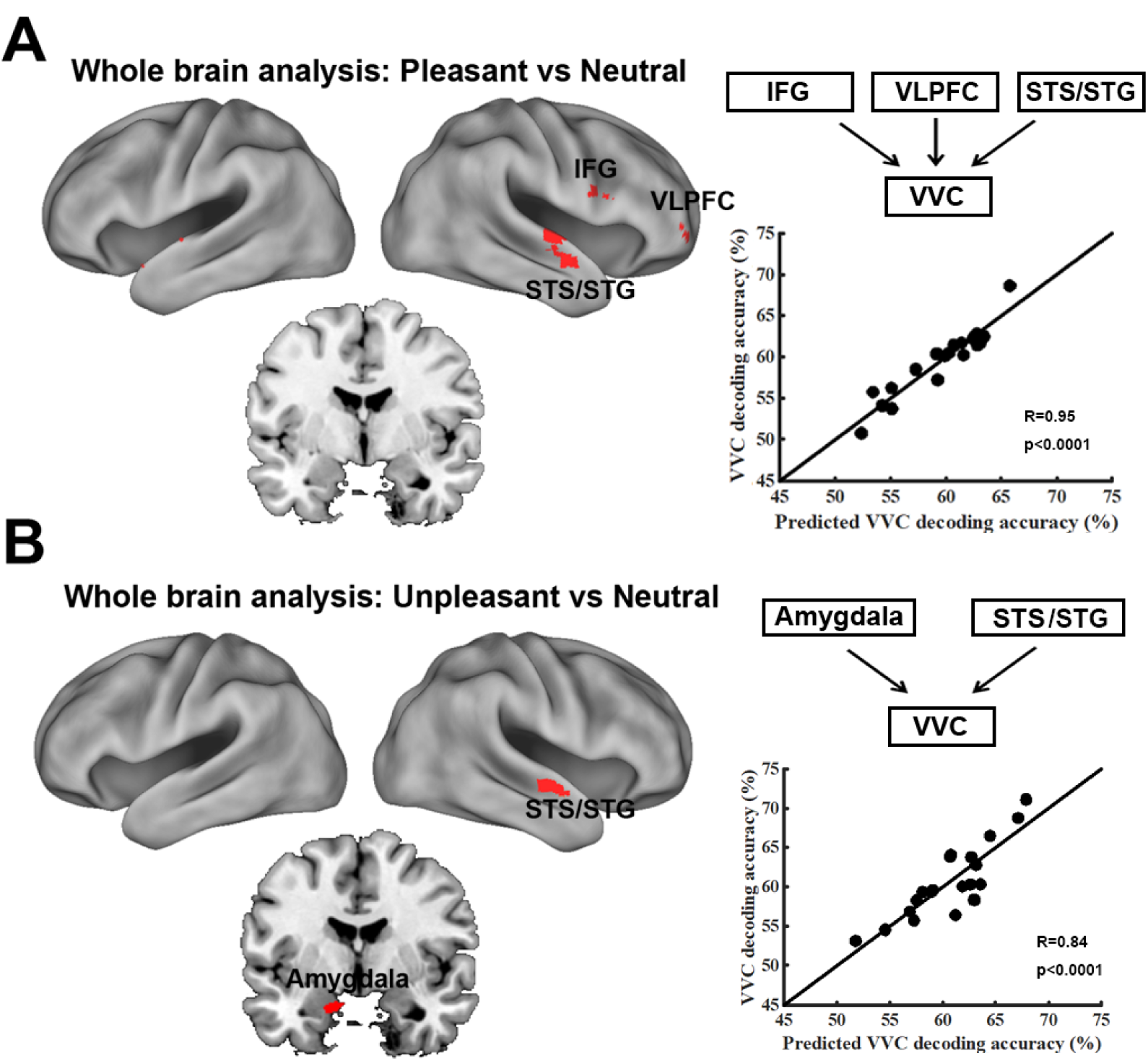
VVC-seeded whole-brain effective connectivity analysis. Effective connectivity from each voxel of the brain onto VVC was computed and correlated with VVC decoding accuracy. Collective contributions of signal reentry from brain regions so identified to VVC decoding accuracy were assessed via multiple regression models. ***A.*** Pleasant vs Neutral (left), where *VVC decoding accuracy* = *β*_0_ + *β*_1_(*IFG* → *VVC*) + *β*_2_ (*VLPFC* → *VVC*) + *β*_3_ (*STS*/*STG* → *VVC*) + ɛ (right). ***B***. Unpleasant vs Neutral (left), where *VVC* decoding accuracy = *β*_0_ + *β*_1_(Amygadala → *VVC*) + *β*_2_(*STS*/*STG* → *VVC*) + ɛ (right). STG, superior temporal gyrus; STS, superior temporal sulcus; IFG, inferior frontal gyrus; VLPFC, ventral lateral prefrontal cortex. All maps were thresholded at R>0.61, p<0.005 and clusters containing more than 10 contiguous such voxels are shown.

A further analysis is conducted to test whether LPP as an index of signal reentry and the frontotemporal effective connectivity to ventral visual cortex is related. Again, a multiple regression model was used with LPP as predicted variable and effective connectivities (EC) as predictor variables. That is, for Pleasant: *LPP* = *β*_0_ + *β*_1_(*IFG* → *VVC*) + *β*_3_(*STS*/*STG* → *VVC*) + ɛ; for Unpleasant,. The predicted LPP and recorded LPP for pleasant pictures and for unpleasant pictures were correlated at R=0.63 (p=0.002) and R=0.56 (p=0.01) respectively. These results affirm prior assertions that LPP is an index of signal reentry and provide further insights into the neural substrate of LPP by identifying the specific frontotemporal sources of these reentrant signals.

## Discussion

We examined the visuocortical representations of affective scenes by recording simultaneous EEG-fMRI from subjects viewing pleasant, unpleasant, and neutral pictures from the IAPS library. Consistent with previous reports, relative to neutral scenes, both pleasant and unpleasant scenes evoked enhanced LPP on the scalp and stronger BOLD activation in a large-scale brain network that included limbic and frontal structures as well as higher-order visual cortices. Applying MVPA to retinotopic visual ROIs, we further found that the multivoxel patterns evoked by pleasant and unpleasant scenes were distinct from those evoked by neutral scenes in all retinotopic visual regions, including the primary visual cortex V1. Concurrently recorded LPP amplitude, an electrophysiological index of recurrent signaling between anterior brain areas and visual cortex, was shown to predict decoding accuracy in ventral visual cortex. This was elaborated by an effective connectivity analysis using DAG, which demonstrated that for unpleasant scenes, the amygdala was a likely source of the reentrant signals, whereas for the pleasant scenes, frontal lobe structures including right IFG and right VLPFC were found to be the likely sources of the reentrant signals.

### Representation of emotional scenes in retinotopic visual cortex

Standardized emotional pictures have been extensively used in human neuroimaging research (see Sabatinelli et al., 2011 for a review), and the resulting body of work has strongly supported the notion that heightened visuocortical engagement is an integral part of an observer’s emotional response. The canonical network selectively activated by affective scenes includes anterior temporal lobe, limbic structures, and prefrontal cortex (Sabatinelli et al., 2011). Mixed findings, however, have been reported regarding the involvement of retinotopic visual areas in the representation of emotional content: Many electrophysiological studies have shown differential retinotopic responses to emotional versus neutral visual cues, both in human observers (Keil et al., 2003; Thigpen et al., 2017) as well as in experimental animals (Li et al., 2019). Similarly, the existence of emotion-specific signals in retinotopic areas is predicted by theoretical work based on animal model data (Amaral et al., 2003), as well as on computational models (Kragel et al., 2019). By contrast, univariate BOLD analyses of retinotopic ROIs, including all early visual areas, tend to not show differences in activation as a function of emotional content (Sabatinelli et al., 2009; meta-analysis in Sabatinelli et al., 2011). This well-established finding was also replicated in the present report, where retinotopic visual areas as well as some higher order visual cortices like PHC1, PHC2 and V3b, were found to be not differentially activated by emotional pictures. Only after we applied multivariate techniques did we find strong evidence for content-specific visuocortical engagement, consistent across a wide range of retinotopic visual regions including primary visual cortex V1.

### Multivariate versus univariate methods

To date, much of fMRI-based neuroscience research has relied on univariate analysis techniques (e.g., GLM analysis). In univariate fMRI analysis, for a voxel to be reported as activated by an experimental condition, it needs to be consistently activated across individuals. As such, individual differences in voxel activation patterns could lead to failure to detect the presence of neural activity in a given region of the brain. Multivariate pattern analysis (MVPA) overcomes this limitation. In MVPA, multivoxel patterns of activation within a ROI are the unit of analysis, and pattern differences between experimental conditions are assessed at the individual subject level, followed by summary statistics computed at the population level (e.g., decoding accuracy averaged across participants). In the past decade, studies have begun to apply MVPA to paradigms where emotional stimuli were used (Said et al., 2010; Kotz et al., 2013; Sitaram et al., 2011; Baucom et al., 2012; Saarimäki et al., 2015; see Kragel and LaBar, 2014 for review). Our study extends this line of research by applying MVPA to study the neural patterns of activation in retinotopic visual areas during affective picture viewing. In particular, when multivoxel patterns of neural activation between emotion and neutral pictures were compared, all retinotopic visual cortices, including primary visual cortex, were shown to contain affective signals, as evidenced by above-chance level decoding accuracies in these regions. Considering that emotional pictures differ from neutral pictures in both emotional arousal and hedonic valence, and these discriminant activation patterns may well reflect differences in either of these two dimensions, we further tested the role of valence in these representations. Decoding between unpleasant and pleasant viewing conditions, in which emotional arousal ratings were largely matched, it was found that the decoding accuracy was above-chance level in nearly all the visual ROIs, suggesting that valence-specific information is encoded in retinotopic visual cortex. This result is consistent with previous work showing that affective valence modulates the gating of early visual input and scope of sensory encoding (Schmitz et al., 2009).

Although the IAPS pictures used in this study were carefully selected for matching content and image composition, an obvious question concerns whether the differences in multivoxel patterns between different emotional categories are attributable to differences in physical properties that are of a non-emotional nature. To address this concern, we conducted a control analysis by subdividing neutral pictures into pictures with neutral people and pictures with neutral scenes. As shown in Figure S1 of the Supplementary Materials, decoding accuracy between neutral people and neutral scenes was at chance level within all retinotopic visual regions, suggesting that it is the emotional content of the pictures rather than the physical/categorical composition (e. g., people versus scenes) that determined the observed pattern differences in the retinotopic regions considered in this study.

### Role of reentrant signals in visual representations of emotion

Where do affective signals in retinotopic visual cortex come from? One recent hypothesis stresses that visual cortex can innately discriminates stimuli varying in affective significance (see Miskovic & Anderson, 2018 for a review). For example, both animal model studies and human studies (Li et al., 2019; Weinberger, 2004; Thigpen et al., 2017) have found that sensory cortex is able to encode the threat content of a visual stimulus, typically after extensive learning. A recent computational study supported this hypothesis by showing that an artificial deep neural network, whose training requires a large amount of stimuli with repetition, has the ability to encode emotional content (Kragel et al., 2019). In human observers, such differential sensitivity to visual features associated with emotional content may be acquired through daily experience (McTeague et al., 2018). However, given that the exemplars used in the present study were novel for the participants, and presented only five times across the duration of the study, it is unlikely that retinotopic visual cortex learned to represent individual features that are linked to emotional significance during the course of the experimental session. Thus, we examined the evidence for the alternative hypothesis that affective signals in retinotopic areas emanate from anterior frontotemporal structures via the mechanism of reentry.

The signal reentry hypothesis proposes that when viewing emotional stimuli, subcortical structures such as the amygdala, upon receiving the initial input, send feedback signals into visual cortex to enhance the processing of the motivationally salient visual input (Sabatinelli et al., 2009; Keil et al., 2009; Lang and Bradley, 2010). Such enhanced visual processing prompts increased vigilance towards appetitive or aversive stimuli, ultimately promoting the deployment of adaptive action in the interest of survival. The reentry hypothesis is indirectly supported by neuroanatomy studies showing feedback projections from the amygdala to the ventral visual stream (Amaral et al., 2003; Freese and Amaral et al., 2005). Patients with amygdala-lesion tend to show no visual cortex enhancement in response to threat, even when their visual cortex is structurally intact (Vuilleumier et al., 2004). In human imaging studies, it has been shown that BOLD activation in amygdala precedes activation in visual cortex, and the functional connectivity between amygdala and ventral visual cortex is increased during affective picture viewing (Sabatinelli et al., 2009; Sabatinelli et al., 2005). Further analysis using Granger causality demonstrated heightened directional connectivity from amygdala to fusiform cortex when viewing emotional scenes (Sabatinelli et al., 2014; Frank et al., 2019).

To examine how signal reentry is affecting neural representations of affective pictures in retinotopic visual cortex, we first used the late positive potential (LPP) to index reentrant processing (Lang et al., 1998; Hajcak et al., 2006; Lang and Bradley, 2010), which was well motivated by prior studies which, combining EEG and fMRI, found covariation between LPP and BOLD activity in multiple brain regions including visual cortices and subcortical structures such as amygdala (Sabatinelli et al., 2012; Liu et al., 2012). Dividing visual cortex into early, ventral, and dorsal portions (referred to as EVC, VVC, and DVC, respectively), we found that for both unpleasant and pleasant pictures, LPP amplitude was statistically significantly correlated with decoding accuracy in VCC, but not in EVC and DVC. This pattern of results is understandable from an anatomical perspective because VVC, beginning at the anterior edge of V4 and extending anteriorly to posterior parahippocampal cortex, receives extensive input from anterior temporal and frontal structures, and is thus expected to be strongly influenced by these structures. Upon receiving reentrant feedback VVC may also play the role of transmitting the reentrant signals down to early visual cortex. Functionally, given that VVC is involved in object and scene perception, which is essential for the current experimental paradigm requiring participants to perceive static pictures with complex affective contents, the purpose of reentry could be to selectively enhance the visual regions mostly related to the ongoing task.

DVC contains higher order visual cortex IPS along the dorsal-parietal pathway. In our data, decoding accuracy in DVC was higher relative to VVC and EVC, but was not correlated with LPP amplitude, suggesting that neural representations of emotional content in DVC was not influenced by reentry. Neuroscientifically, DVC is known to play essential roles in visual spatial attention, motion perception, and motor preparation (Bressler et al., 2008; Wandell et al., 2011). A previous study using naturalistic emotional videos (Goldberg et al., 2014) reported a preferential activation in the dorsal-parietal visual stream, which, the authors hypothesized, was related to motor preparation associated with emotionally salient information. Our results can be seen as lending support to this hypothesis by showing that viewing affective stimuli evokes emotion-specific neural patterns in dorsal association cortices that lie at the interface between visual perception and motor preparation for survival-relevant action (Lang and Bradley, 2010). Furthermore, along with the findings in ventral visual cortex, our results also suggest that viewing emotional scenes prompts parallel emotional processes along ventral and dorsal visual pathways, serving different functional roles.

In addition to LPP amplitude, we also used inter-regional effective connectivity as an index of reentry, with an initial emphasis on the role of amygdala in generating the feedback signals. Correlating VVC decoding accuracy with amygdala→VVC effective connectivity, we found that for unpleasant scenes, decoding accuracy in VVC is associated with amygdala→VVC effective connectivity, lending credence to the idea that the reentrant signals arise from limbic structures and reach visual cortex to modulate the visuocortical representations of emotion. For pleasant scenes, however, amygdala→VVC effective connectivity did not predict VVC decoding accuracy, in contrast to expectations. We then carried out a seed-based whole-brain effective connectivity analysis with VVC as the seed. This analysis revealed that effective connectivity from bilateral STS/STG, right IFG, and right VLPFC influenced decoding accuracy in VVC between pleasant scenes and neutral scenes. Interpreting the finding, we note that VLPFC has been considered part of the ventral affective system which includes VLPFC, medial prefrontal cortex (MPFC), and amygdala (Dolcos and McCarthy, 2006). Anatomically, VLPFC shows strong connection with sensory areas as well as reward processing structures such as amygdala and orbitofrontal cortex (Carmichael and Price, 1995a; Carmichael and Price, 1995b; Petrides and Pandya, 2002; Kennerley and Wallis, 2009). It integrates reward information and provides top-down signals to sensory cortex to improve behavioral performance (Kennerley and Wallis, 2009). Right IFG is known to be involved both in cognitive control and in emotional processing by exerting top-down control of STS/STG (Frye et al., 2010) and ventral sensory stream (Tops and Boksem, 2011). It is worth stressing that our results, for the first time, suggest a role of right IFG and VLPFC in generating feedback signals to influence the sensory processing of appetitive input.

For unpleasant scenes, the whole brain effective connectivity analysis again revealed that the amygdala was an important source of reentrant feedback, along with STS/STG. Whereas the appearance of amygdala in the whole brain map agrees with and confirms the ROI based analysis, the appearance of the STS/STG region in the brain map was a new finding; the same STS/STG region appeared both in the processing of pleasant and unpleasant scenes (Narumoto et al., 2001), suggesting that the STS/STG region may serve a way station which transmits the signals received from the limbic and/or prefrontal structures to visual cortex. Primate studies have found that STS/STG reciprocally interact with the ventral visual stream (Cusick, 1997; Allison et al., 2000). In addition, STS/STG is structurally connected with amygdala and receives its feedback projection (Grezes et al., 2014; Pitcher et al., 2017). As such, the present study helps advance an interesting hypothesis for further studies to test, which, by integrating animal model work, computational modeling, and advanced neuroimaging in humans, can potentially lead to more precisely defined pathways of neural feedback signaling in emotional scene perception.

In this work both LPP and effective connective were used to index signal reentry from frontotemporal cortices. A multiple regression analysis was performed to examine whether they are related. We found that for pleasant picture perception, a large portion of LPP variance was explained by IFG→VVC, VLPFC→VVC, and STS/STG→VVC effective connectives, whereas for unpleasant picture perception, a large portion of LPP variance was explained by amygdala→VVC and STS/STG→VVC effective connectivities. These findings, besides demonstrating that the two measures of signal reentry are related, provide a connectivity level neural substrate of LPP, which has been mainly linked to distributed brain regions without regards to their functional interactions (Sabatinelli et al., 2012; Liu et al., 2012).

### Summary

We recorded simultaneous EEG-fMRI from participant viewing natural images containing affective scenes. Applying multivariate pattern analysis to fMRI data, we found the existence of affective signals in the entire retinotopic visual cortex, including V1. Using scalp potentials as indices of recurrent signaling between anterior brain regions and visual cortex, we found that the emotion-sensitive neural representations in the ventral portion of retinotopic visual cortex were related to signal reentry from anterior brain structures. Further analysis using effective connectivity further identified the sources of re-entrant signals, which include amygdala and STS/STG in the case of unpleasant scene processing, and IFG, VLPFC and STS/STG in the case of pleasant scene processing.

## Materials and Methods

### Participants

The experimental protocol was approved by the Institutional Review Board of the University of Florida. Twenty-six healthy volunteers with normal or corrected-to-normal vision gave written informed consent and participated in this study. Before the MRI scan, two subjects withdrew from the experiment. In addition, data from four participants were discarded due to artifacts generated by excessive movements inside the scanner. The data from the remaining twenty subjects were analyzed and reported here (10 women; mean age: 20.4±3.1).

### Stimuli

The stimuli consisted of 60 pictures selected from the International Affective Picture System (IAPS) (Lang et al., 1997). Their IAPS IDs are: (1) 20 pleasant pictures: 4311, 4599, 4610, 4624, 4626, 4641, 4658, 4680, 4694, 4695, 2057, 2332, 2345, 8186, 8250, 2655, 4597, 4668, 4693, 8030; (2) 20 neutral pictures: 2398, 2032, 2036, 2037, 2102, 2191, 2305, 2374, 2377, 2411, 2499, 2635, 2347, 5600, 5700, 5781, 5814, 5900, 8034, 2387; (3) 20 unpleasant pictures: 1114, 1120, 1205, 1220, 1271, 1300, 1302, 1931, 3030, 3051, 3150, 6230, 6550, 9008, 9181, 9253, 9420, 9571, 3000, 3069. The pleasant pictures included sport scenes, romance, and erotic couples, and had an average valence rating of 7.0 ± 0.45. The unpleasant pictures included threat/attack scenes and bodily mutilations and had an average valence rating of 2.8 ± 0.88. The neutral pictures consisted of images containing landscapes and neutral humans and had an average valence rating of 6.3 ± 0.99. The arousal ratings of these pictures are as follows: pleasant (5.8 ± 0.90), unpleasant (6.2 ± 0.79), both being higher than that of neutral pictures (4.2 ± 0.97). (All ratings are based on a 1-9 scale.) To minimize confounds, across categories, pictures should be similar overall in composition and in rated complexity and matched in picture file size.

### Procedure

Each IAPS picture was displayed on a MR-compatible monitor for 3s, which was followed by a variable (2800 ms or 4300 ms) interstimulus interval. There were 5 sessions. Each session comprised 60 trials corresponding to the 60 different pictures. The same 60 pictures were shown in each session, but the order of picture presentation was randomized from session to session. Stimuli were presented on the monitor placed outside the scanner and viewed via a reflective mirror. Participants were instructed to maintain fixation on cross at the center of the screen during the whole session. After the experiment, participants were instructed to rate the hedonic valence and emotional arousal level of 12 representative IAPS pictures (4 pleasant, 4 neutral and 4 unpleasant) which were not part of the 60-picture set. The rating was done using a paper and pencil version of the self-assessment manikin (Bradley and Lang, 1994). As shown in Table S2, the ratings of the 12 pictures by the participants are consistent with the normative ratings of these pictures (Lang et al., 1997).

### Data acquisition

Functional MRI data were collected on a 3T Philips Achieva scanner (Philips Medical Systems), with the following parameters: echo time (TE), 30 ms; repetition time (TR), 1.98 s; flip angle, 80°; slice number, 36; field of view, 224 mm; voxel size, 3.5*3.5*3.5 mm; matrix size, 64*64]. Slices were acquired in ascending order and oriented parallel to the plane connecting the anterior and posterior commissure. T1-weighted high-resolution structural image was also obtained.

EEG data were recorded simultaneously with fMRI using a 32 channel MR-compatible EEG system (Brain Products GmbH). Thirty-one sintered Ag/AgCl electrodes were placed on the scalp according to the 10-20 system, and one additional electrode was placed on subject’s upper back to monitor electrocardiograms (ECGs). The recorded ECG was used to detect heartbeat events and assist in the removal of the cardioballistic artifacts. The EEG channels were referenced to the FCz electrode during recording. EEG signal was recorded with an online 0.1-250Hz band-pass filter and digitized to 16-bit at a sampling rate of 5 kHz. The EEG recording system was synchronized with the scanner’s internal clock throughout recording to ensure the successful removal of the gradient artifact in subsequent preprocessing.

### Data preprocessing

The fMRI data were preprocessed with SPM (http://www.fil.ion.ucl.ac.uk/spm/). The first five volumes from each session were discarded to eliminate artifacts caused by the transient instability of scanner. Slice timing was corrected using interpolation to account for differences in slice acquisition time. The images were then corrected for head movement by spatially realigning them to the sixth image of each session, normalized and registered to the Montreal Neurological Institute (MNI) template, and resampled to a spatial resolution of 3×3×3 mm. The transformed images were smoothed by a Gaussian filter with a full width at half maximum of 8 mm. Low frequency temporal drift was removed from the functional images by applying a high-pass filter with a cutoff frequency of 1/128 Hz.

The EEG data were first processed using Brain Vision Analyzer 2.0 (Brain Products GmbH) to remove scanner artifacts. For removing gradient artifacts, a modified version of the original algorithm proposed for this purpose was applied (Allen et al., 2000). Briefly, an artifact template was created by segmenting and averaging the data according to the onset of each volume and subtracted from the raw EEG data. The cardioballistic artifact was removed by an average artifact subtraction method (Allen et al., 1998). Specifically, R peaks were detected in the low-pass-filtered ECG signal and used to establish a delayed average artifact template over 21 consecutive heartbeat events by sliding-window approach. The artifact was then subtracted from the original EEG signal. After gradient and cardioballistic artifacts were removed, the EEG data were low-pass filtered with the cutoff set at 50 Hz, downsampled to 250 Hz, re-referenced to the average reference, and exported to EEGLAB for further processing (Delorme and Makeig, 2004). Second-order blind identification (SOBI) (Belouchrani et al., 1993) was then applied to correct for eye blinking, residual cardioballistic and movement-related artifacts. The artifact-corrected data were epoched from −300ms to 2000ms with 0ms denoting picture display onset. The prestimulus baseline was defined as −300ms to 0ms for ERP analysis.

### Single-trial estimation of fMRI-BOLD

Trial-by-trial picture-evoked BOLD signal was estimated by using the beta series method (Mumford et al., 2012). In this method, the trial of interest was represented by one regressor and all the other trials were represented by another regressor. Six motion regressors were also included to account for any movement-related artifacts during scan. Repeating the process for all the trials we obtained the BOLD response to each picture presentation in all brain voxels. These single trial BOLD responses were used in the MVPA decoding analysis.

### Regions of interest in retinotopic visual cortex

Regions of interest (ROIs) were defined according to a recently published probabilistic visual retinotopic atlas (Wang et al. 2014). Seventeen ROIs in the retinotopic atlas were included: V1v, V1d, V2v, V2d, V3v, V3d, V3a, V3b, hV4, VO1, VO2, PHC1, PHC2, LO1, LO2, hMT, and IPS; see Figure 3A. The homologous regions from the two hemispheres were combined, and the IPS ROI was formed by combining the voxels from IPS0, IPS1, IPS2, IPS3, IPS4, and IPS5.

### Multivariate pattern analysis

MVPA was performed by the linear support vector machine (SVM) method using the LibSVM package (http://www.csie.ntu.edu.tw/~cjlin/libsvm/) (Chang and Lin, 2011). Single-trial voxel patterns evoked by pleasant, unpleasant, and neutral pictures were decoded in a pairwise fashion (e.g., pleasant vs neutral) within 17 retinotopic ROIs. The classification accuracy was calculated by ten-fold validation. Specifically, all the data was divided into ten equal sub-datasets, nine of which comprised training data to train the classifier and the remaining one of which was used for testing. The decoding accuracies of ten such procedures were averaged. To further ensure the stability of the decoding result, we repeated ten-fold partition one hundred times, conducted the same procedure to acquire decoding accuracies for each partition, and averaged the accuracies to yield the decoding accuracy for each ROI. Repeating the process for each participant, group level decoding accuracy was computed by averaging individual accuracies across twenty participants. A non-parametric permutation-based technique (Stelzer et al. 2013) was applied to test the whether the statistical significance of decoding accuracy was above chance level for each ROI. At the individual subject level, the class labels were randomly shuffled 100 times and each shuffled run generated one chance level decoding accuracy. At the group level, one of the chance level decoding accuracies was extracted randomly from each subject and averaged across subjects. This process was repeated 105 times to obtain an empirical distribution of the chance level accuracy at the group level for each ROI. The accuracy corresponding to p=0.001 in the probability distribution of the permutation test was used as the threshold to determine whether decoding accuracy is above chance level.

### Effective connectivity via directed acyclic graphs (DAG)

Functional connectivity analysis is typically based on cross correlation analyses. Cross correlation has a key limitation: it does not provide directional information. To test the potential sources of reentrant signals, we thus used an effective connectivity measurement called the directed acyclic graph (DAG), derived from linear non-Gaussian acyclic models (LiNGAM) (Shimizu et al.,2006; Shimizu et al.,2011). LiNGAM is a linear non-Gaussian variant of structural equation modeling (SEM). It has been successfully applied in fMRI work to assess effective connectivity (Schlösser et al., 2003; Marrelec et al., 2009; Liu et al., 2015). Briefly, LiNGAM estimates causal order using the recurrent regression method to find exogenous variables in the system and analyze the connection strength via least squares regression. The algorithm achieves better efficiency when prior knowledge is provided. In the current work, given the evidence from prior anatomical and causality studies which showed direct synaptic connection and significantly greater directional connection from amygdala to the ventral visual system (Amaral et al., 2003; Sabatinelli et al.,2014), we provided the prior knowledge that signal in amygdala leads the directional connection towards VVC. With such prior knowledge, the estimation of LiNGAM is equivalent to a structural equation model. The connection coefficient from LiNGAM is used to represent effective connectivity strength.

Two types of effective connectivity analysis were considered: one based on *a priori* considerations and the other a seed-based whole-brain analysis. In the analysis based on the *a priori* considerations, ROIs were the amygdala and the ventral visual cortex. Amygdala ROI consisted of two spherical masks of 5 mm in radius centered at [–16, 0, −24] and [20, 0, −20] (Hamann et al., 2004). The ventral visual cortex (VVC) ROI consisted of retinotopic visual regions VO1, VO2, PHC1, and PHC2. For the seed-based whole-brain analysis, VVC was used as the seed, and the effective connectivity into VVC was assessed for all the voxels in the brain.

### ERP analysis

The late positive potential (LPP) was estimated at electrode Pz. Preprocessed EEG data (see above) was low-pass filtered at 30Hz and averaged within each picture category to yield the event-related potential for that category. In addition to grand average LPP, the LPP amplitude for each subject was obtained using the time interval of 500 ms in duration around the peak of LPP (Liu et al., 2012).

## Acknowledgements

This work was supported by NIH grants R01 MH112558.

## Supplementary Materials

In the supplementary material section, we address three issues. The first issue concerns whether the differences in multivoxel patterns between different emotional categories are attributable to differences in non-emotional content. We divided the neutral pictures into two subcategories, neutral people and adventure/nature scene, according to the categorical label from IAPS. These two subcategories are affectively neutral but vary in non-emotional content. Pictures of neutral people contain human body and face whereas pictures of adventure/nature scenes are landscapes and the like. We conducted the same decoding analysis as in Figure 3. The decoding accuracy in each retinotopic ROI is shown in Figure S1. Group level random permutation test was conducted, and the statistically significant threshold was determined to be 55.5%, which corresponded to p=0.001. No region has above chance decoding.

The second issue concerns the abbreviations used in the main body of the manuscript. A table of explanations of these abbreviations are provided (Table 1).

The third issue concerns the consistency between the ratings of the 12 IAPS pictures by the participants of this study and the corresponding normative ratings of the same pictures. Following scanning, participants rated hedonic valence and emotional arousal level of 12 representative IAPS pictures (4 pleasant, 4 neutral and 4 unpleasant) which were not part of the 60-picture set. The results showed that the valence and arousal ratings by the participants were consistent with the normative ratings (Table S2).

**Figure S1.**
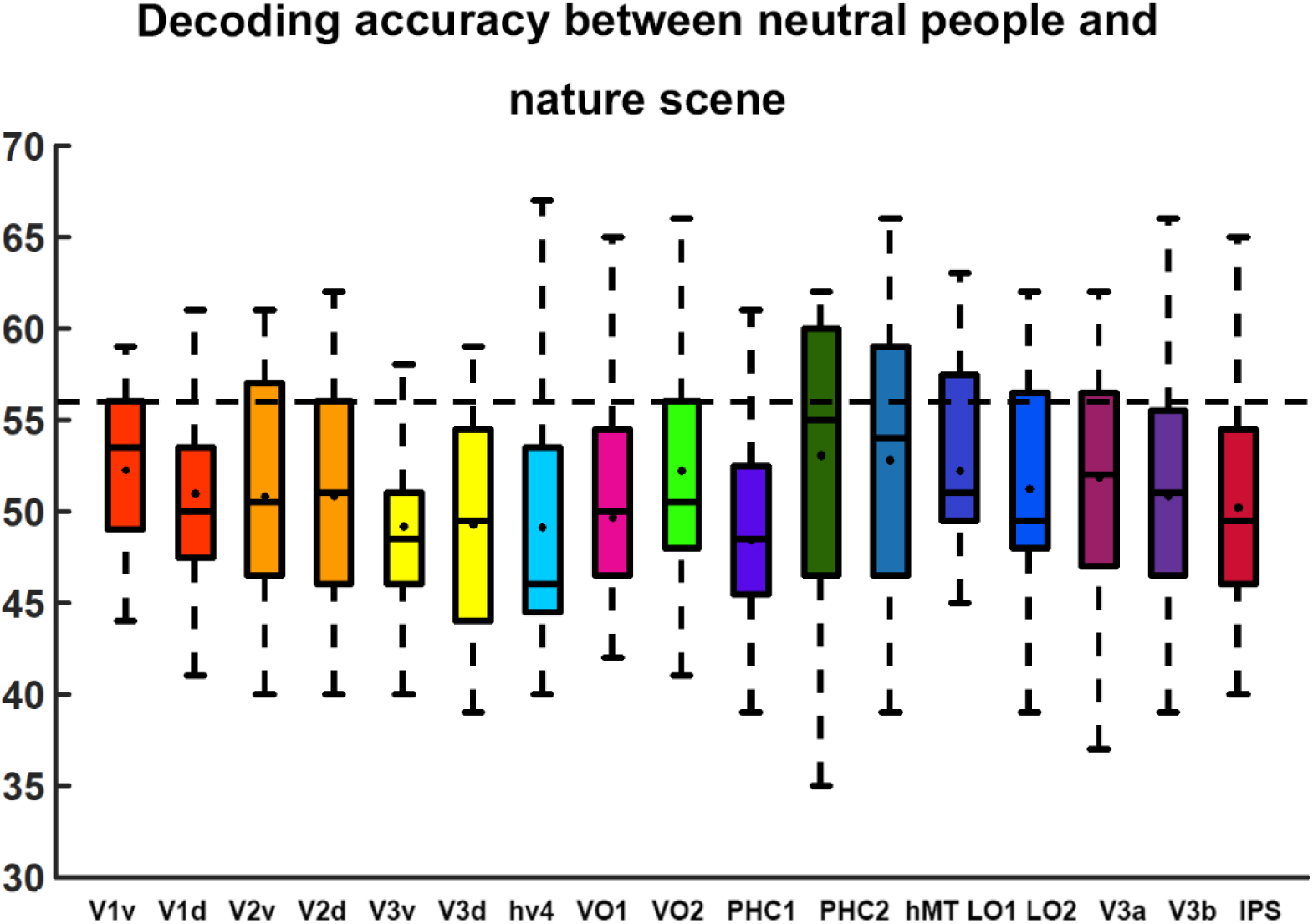
Decoding accuracy between neutral people versus nature scenes SVM across all retinotopic ROIs. All decoding accuracies are not significantly different from chance level of 50%. Statistical threshold is at 55.5% (p=0.001) according to a random permutation test.

**Table S1.**
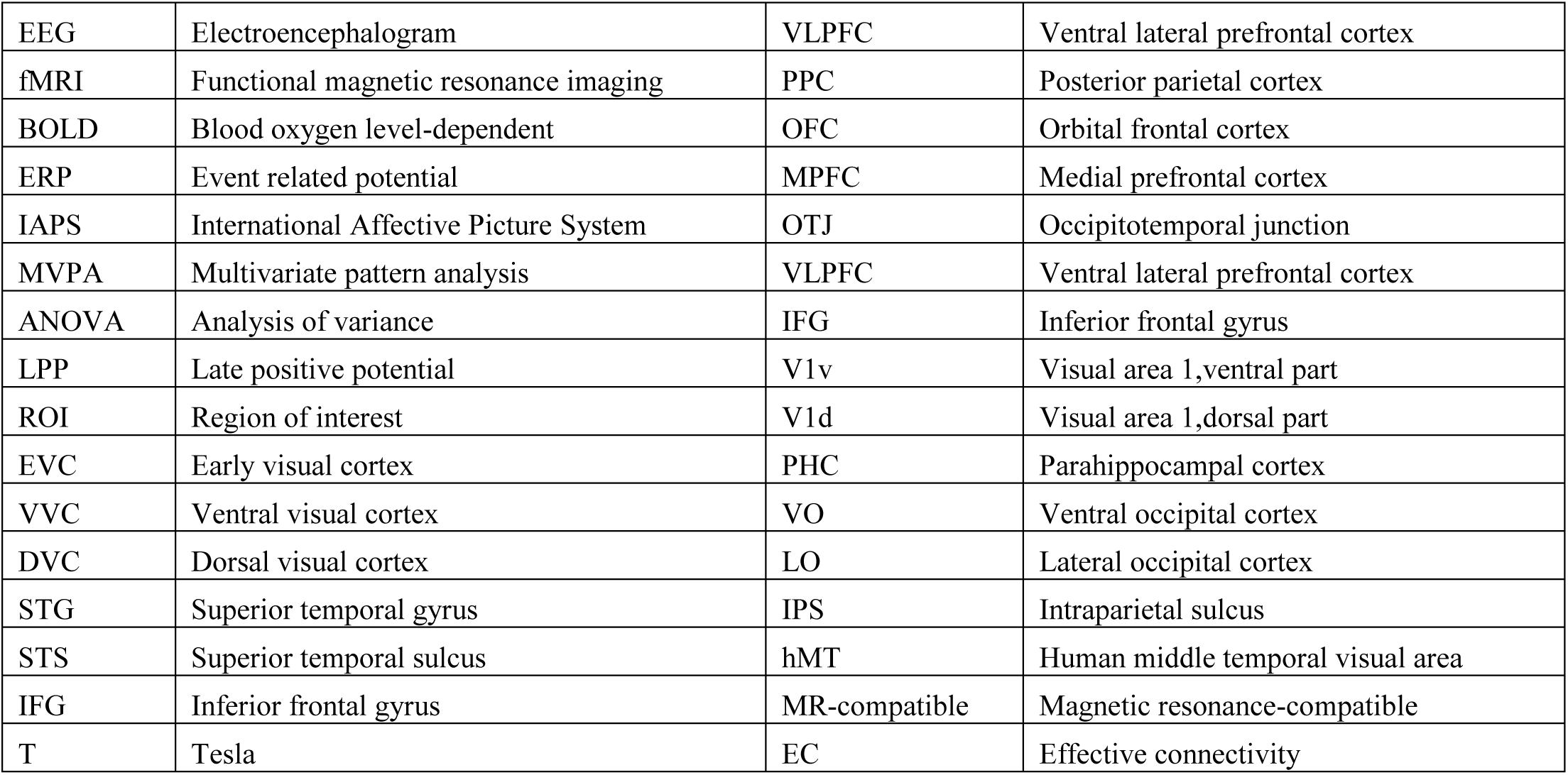
Full expressions for the abbreviations used in the text.

**Table S2.**
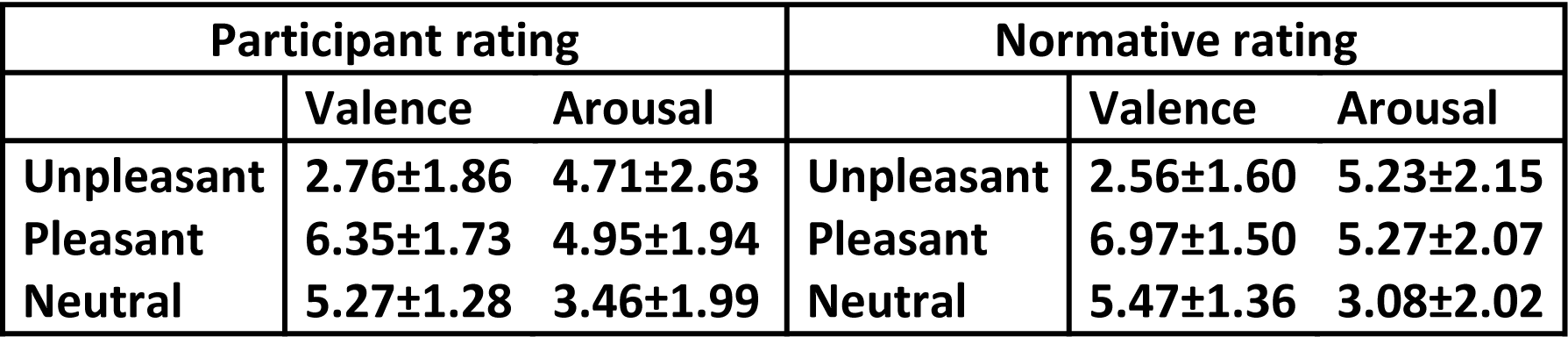
Ratings of the 12 IAPS images by the participants and the normative ratings of the same pictures. No statistically significance difference was found between the two sets of ratings.

