## Supplementary material for "Decoding Multivoxel Representations of Affective Scenes in Retinotopic Visual Cortex": Supplementary Materials final.docx

In the supplementary material section, we address three issues. The first issue concerns whether the differences in multivoxel patterns between different emotional categories are attributable to differences in non-emotional content. We divided the neutral pictures into two subcategories, neutral people and adventure/nature scene, according to the categorical label from IAPS. These two subcategories are affectively neutral but vary in non-emotional content. Pictures of neutral people contain human body and face whereas pictures of adventure/nature scenes are landscapes and the like. We conducted the same decoding analysis as in Figure 3. The decoding accuracy in each retinotopic ROI is shown in Figure S1. Group level random permutation test was conducted, and the statistically significant threshold was determined to be 55.5%, which corresponded to p=0.001. No region has above chance decoding.

The second issue concerns the abbreviations used in the main body of the manuscript. A table of explanations of these abbreviations are provided (Table 1).

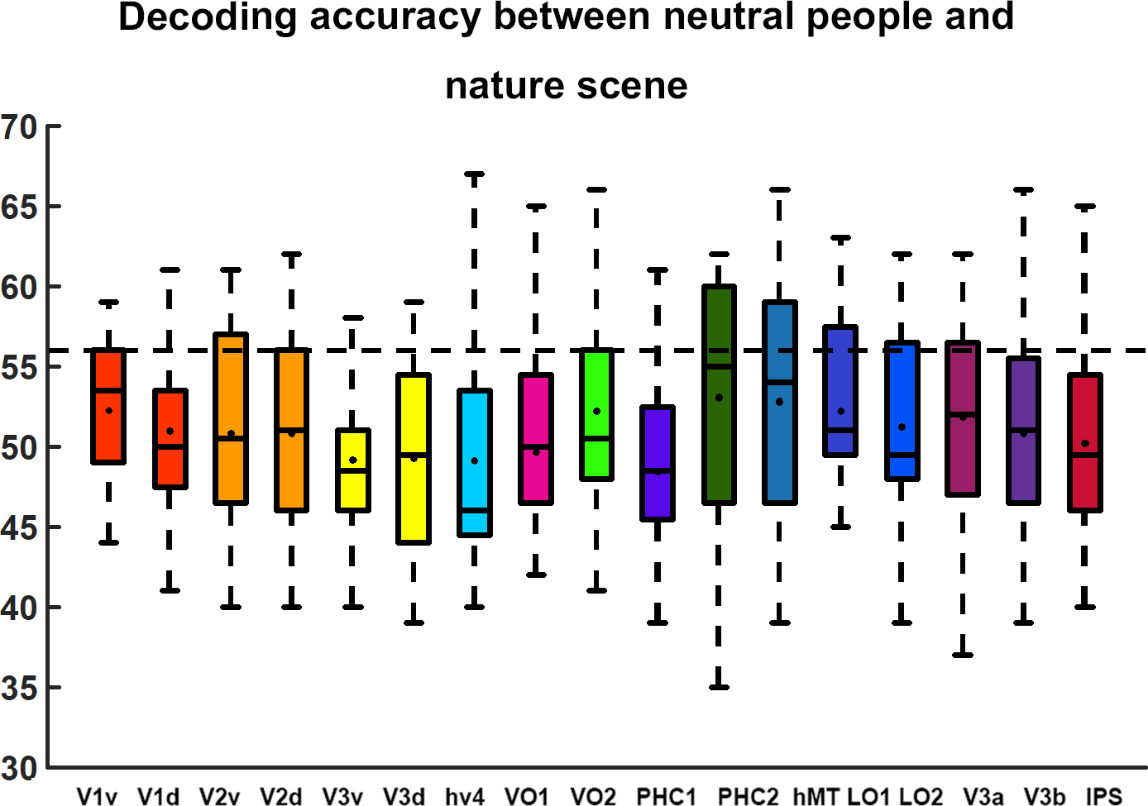
The third issue concerns the consistency between the ratings of the 12 IAPS pictures by the participants of this study and the corresponding normative ratings of the same pictures. Following scanning, participants rated hedonic valence and emotional arousal level of 12 representative IAPS pictures (4 pleasant, 4 neutral and 4 unpleasant) which were not part of the 60-picture set. The results showed that the valence and arousal ratings by the participants were consistent with the normative ratings (Table S2).

Figure S1. Decoding accuracy between neutral people versus nature scenes SVM across all retinotopic ROIs. All decoding accuracies are not significantly different from chance level of 50%. Statistical threshold is at 55.5% (p=0.001) according to a random permutation test.

| EEG | Electroencephalogram | VLPFC | Ventral lateral prefrontal cortex |
| --- | --- | --- | --- |
| fMRI | Functional magnetic resonance imaging | PPC | Posterior parietal cortex |
| BOLD | Blood oxygen level-dependent | OFC | Orbital frontal cortex |
| ERP | Event related potential | MPFC | Medial prefrontal cortex |
| IAPS | International Affective Picture System | OTJ | Occipitotemporal junction |
| MVPA | Multivariate pattern analysis | VLPFC | Ventral lateral prefrontal cortex |
| ANOVA | Analysis of variance | IFG | Inferior frontal gyrus |
| LPP | Late positive potential | V1v | Visual area 1,ventral part |
| ROI | Region of interest | V1d | Visual area 1,dorsal part |
| EVC | Early visual cortex | PHC | Parahippocampal cortex |
| VVC | Ventral visual cortex | VO | Ventral occipital cortex |
| DVC | Dorsal visual cortex | LO | Lateral occipital cortex |
| STG | Superior temporal gyrus | IPS | Intraparietal sulcus |
| STS | Superior temporal sulcus | hMT | Human middle temporal visual area |
| IFG | Inferior frontal gyrus | MR-compatible | Magnetic resonance-compatible |
| T | Tesla | EC | Effective connectivity |

Table S1. Full expressions for the abbreviations used in the text.

| **Participant rating** | | | **Normative rating** | | |
| --- | --- | --- | --- | --- | --- |
|  | **Valence** | **Arousal** |  | **Valence** | **Arousal** |
| **Unpleasant** | **2.76±1.86** | **4.71±2.63** | **Unpleasant** | **2.56±1.60** | **5.23±2.15** |
| **Pleasant** | **6.35±1.73** | **4.95±1.94** | **Pleasant** | **6.97±1.50** | **5.27±2.07** |
| **Neutral** | **5.27±1.28** | **3.46±1.99** | **Neutral** | **5.47±1.36** | **3.08±2.02** |

Table S2. Ratings of the 12 IAPS images by the participants and the normative ratings of the same pictures. No statistically significance difference was found between the two sets of ratings.
